# Extent of adaptation is not limited by unpredictability of the environment in laboratory populations of *Escherichia coli*

**DOI:** 10.1101/165696

**Authors:** Shraddha Karve, Devika Bhave, Sutirth Dey

## Abstract

Environmental variability is on the rise in different parts of the earth and the survival of many species depend on how well they cope with these fluctuations. Our current understanding of how organisms adapt to unpredictably fluctuating environments is almost entirely based on studies that investigate fluctuations among different values of a single environmental stressor like temperature or pH. However, in nature multiple stresses often exist simultaneously. How would unpredictability in environmental fluctuations affect adaptation under such a scenario? To answer this question, we subjected laboratory populations of *Escherichia coli* to selection over ~260 generations. The populations faced predictable and unpredictable environmental fluctuations across qualitatively different selection environments, namely, salt and acidic pH. We show that predictability of environmental fluctuations does not play a role in determining the extent of adaptation. Interestingly, the extent of ancestral adaptation, to the chosen selection environments, is of key importance. Integrating the insights from two previous studies, our results suggest that it is the simultaneous presence of multiple environmental factors that poses a bigger constraint on extent of adaptation, rather than unpredictability of the fluctuations.

## Introduction

Increasing climatic variability around the globe [1, 2] has exposed many natural populations to greater environmental fluctuations than what they have experienced in their recent evolutionary past. Since the ability of these organisms to face environmental fluctuations will determine the survival of the species, over the last decade or so, a large number of studies have investigated the ecological and evolutionary response of organisms to such fluctuations [3-7]. These studies have shown that, among other things, the frequency and predictability of the environmental fluctuations can affect the fitness outcomes. In asexual organisms, for instance, phenotypic plasticity can be expected to evolve when environment changes rapidly i.e. within few generations [8]. On the other hand, where environment changes after few hundred generations, large effects mutations are expected to sweep through the population [9, 10].

Similar predictions have been made and tested regarding the fitness outcomes of predictable and unpredictable fluctuations. For instance, theory suggests that when fluctuations in selection environments are not correlated, mean fitness of the populations is not expected to change [11]. Experiments show that this is indeed the case when environment fluctuates unpredictably over short duration of selection [12]. However, when faced with unpredictable environments over longer duration, the overall mean fitness actually improves [13]. Another theoretical study predicts, and further experimentally validates, that asexual organisms experiencing predictable fluctuations on short time scales would evolve switching phenotypes with switching rates that are tuned to the rate of environmental change [14]. Furthermore, bacterial populations experiencing predictable fluctuations have been shown to improve fitness over all the values of environments experienced during the selection [15-18]. Fitness outcomes of unpredictable fluctuations, on the other hand, are more difficult to predict. Selection in unpredictable fluctuations can lead to increase in fitness in the selection environments [18, 19] or show no change [3]. To complicate matters further, in some cases, fitness can increase for some selection environments but not for others [16] or for some measures of fitness but not for others [5]. Viral populations facing unpredictable fluctuations in host types can improve fitness in both the hosts experienced during the selection [18]. On the other hand, bacterial populations experiencing different values of pH unpredictably, can improve fitness for some of the pH values but not others [16]. Interestingly, viral populations facing stochastic changes in temperature improve fitness at the highest value of temperature experienced during the selection. However, the magnitude of this fitness gain is less compared to the populations facing predictable temperature changes [3]. In short, outcomes of unpredictable fluctuations remain elusive and demands further investigation.

In spite of the lack of consensus on effects of selection under unpredictability, experimental studies in this area have led to several interesting observations. When the selection environment fluctuates between different values of the same parameter (say different values of temperature [3, 17, 19, 20], then although the strength of selection changes, the nature of selection remains similar. In other words, when an organism adapts to a temperature of say 40 ºC, then it is likely to do well under 39 ºC or 38ºC. However, when the environment fluctuates across qualitatively different parameters (say salt and pH [12, 13], then the both the nature and the strength of selection can vary. This change is likely to affect the adaptation of organisms. What would be the effects of unpredictability in environmental fluctuations under such circumstances? Under fluctuating environments, populations might face the extreme magnitudes of a stress intermittently. Does this affect evolutionary adaptation [3]? Specifically, do populations that experience more of the extreme values for a given stress (say temperature) during environmental fluctuations, adapt better to that stress?

In this study, we attempt to answer some of these questions, pertaining to unpredictability, using experimental evolution of laboratory populations of *E. coli* under fluctuating environments. We show that unpredictability does not hinder the extent of adaptation when environment fluctuates across different environmental parameters. In fact, the extent of adaptation, in this case, is mainly governed by how well the ancestor is adapted to a chosen selection environment. Our results suggest that the fitness of the ancestor, in the chosen selection environments, is an important fitness determinant along with the predictability and grain size of the fluctuations. Additionally, at least in our study, the number of instances of exposure to the extreme values of the selection environment do not play a role in determining the adaptation of the populations facing fluctuating environments.

## Methods

### Selection

We used Kanamycin resistant *Escherichia coli* strain K12 (see S1 for details) for the selection experiment. A single colony grown on Nutrient agar with Kanamycin (see SOM for composition) was inoculated in 50 ml of Nutrient broth with Kanamycin (NB^Kan^) (see S2 for composition) and allowed to grow for 24 hr at 37^0^C, 150 rpm. 120 replicate populations were initiated by adding 4 µl of this suspension to 2 ml of NB^Kan^. These 120 replicate populations were randomly assigned to six different selection regimes, 20 replicate populations per regime.

The control populations were sub-cultured in NB^Kan^ for the entire duration of the selection. pH 4.5 and 5g% salt formed the two constant selection regimes. In one of the predictability regimes ((henceforth PBin, for **P**redictable **Bin**ary) the environment alternated between pH 4.5 and salt 5g% every 24 hours. In the second predictability regime (henceforth termed as UpBin, for **U**n**p**redictable **Bin**ary), the populations faced a random sequence of pH 4.5 or salt 5g% (i.e. the magnitude of the two stresses remained the same but the order in which they were presented became unpredictable). In the final predictability regime (henceforth termed as UpRange for **Unp**redictable **Range**), the environment fluctuated unpredictably over a range of values of salt and acidic pH, (i.e. the magnitude of the stress varied stochastically within a range and their order of presentation was unpredictable) (see S3 for details of all the selection regimes).

24 welled plates with 2 ml of appropriate growth medium and 4 µl of inoculum volume for each well were used throughout the selection and assay experiments. The growth conditions were maintained at 37^0^C, 150 rpm and all the populations were sub-cultured every 24 hours. The selection lasted for 30 days which translates into ~ 260 generations [21]. At the end of the selection, populations were stored as glycerol stocks at -80 ^0^C for fitness assays.

### Fitness measurement

We estimated fitness for all the selected populations in pH 4.5, salt 5g% and NB^Kan^ as per previous study [13]. Briefly, 4 µl of culture was revived overnight in 2 ml of NB^Kan^ from every glycerol stock. 4 µl of this revived culture was inoculated in 2ml of relevant assay environment in 24 welled plate. OD600 was measured every 2 h on a plate reader (Synergy HT BioTek, Winooski, VT, USA) for the period of 24 h. For every assay environment two independent trials were conducted, for all the selection regimes, on two different days resulting in total of 720 fitness estimates.

Fitness was estimated as the maximum growth rate over the period of 24 hours [12, 19] and maximum density reached in 24 h, henceforth referred as *K* [13].

### Statistical analysis

a. Overall mean fitness Fitness estimates were normalized using previously computed [13] average fitness value of the ancestor in the same assay environment, over 20 replicate measurements. The normalized fitness values were analyzed using four-way mixed model ANOVA. Selection (six levels: control, pH 4.5, salt, PBin, UpBin, Uprange) and assay environment (three levels: NB^Kan^, pH 4.5 and salt 5g %) were fixed factors. Replication (twenty levels) was a random factor nested in Selection while Trial (two levels) was used as a block. For the significant main effect of Selection, post-hoc analysis was performed using Tukey’s HSD.
b. Variation for fitness Since variance and standard deviation scale with mean, we estimated coefficients of variation (CV) as a measure of variation in fitness. For a given selected population in a given assay environment, we estimated the average fitness across the two trials. For each population in a given selection regime, we then computed the CV across the fitness estimates in the three different assay environments. Every selection regime thus yielded 20 CV estimates. These were then analyzed using one-way ANOVA with Selection (three levels) as a fixed factor, followed by Tukey’s HSD.

To assess the biological significance of the difference in overall mean fitness of selected and control populations, we computed Cohen’s *d* statistic [22] as a measure of the effect sizes [23]. The effect sizes was interpreted as small, medium and large for 0.2 < *d* < 0.5, 0.5 < *d* < 0.8 and *d* > 0.8, respectively [22]. All ANOVAs in this study were performed on STATISTICAv5.0 (Statsoft Inc., Tulsa, OK, USA), whereas the Cohen’s *d* statistics were estimated using the freeware Effect Size Generator v2.3.0 [24].

## Results

Overall mean fitness, when normalized by the ancestral values, was more than one for all the selected populations, except the control populations grown in NB (Fig 1). This suggests that all populations experienced adaptation during the selection. For both measures of fitness, i.e. maximum growth rate and *K*, the overall mean fitness of the control populations was significantly lower than all other selection regimes (Fig 1, *F_5, 342_* = 13.33, *p* < 0.001 for maximum growth rate, *F_5, 342_* = 20.86, *p* < 0.001 for *K*, Table S4 and Table S5). The effect sizes of the fitness differences between control and other selected populations were medium or large in all cases, except for populations selected in pH 4.5 (Table S6). Among the three fluctuating treatments, overall mean fitness was comparable for both measurements of fitness (Fig 1, Table S5). This suggested that unpredictability posed little hindrance to the adaptation during the adaptation.

**Fig 1.**
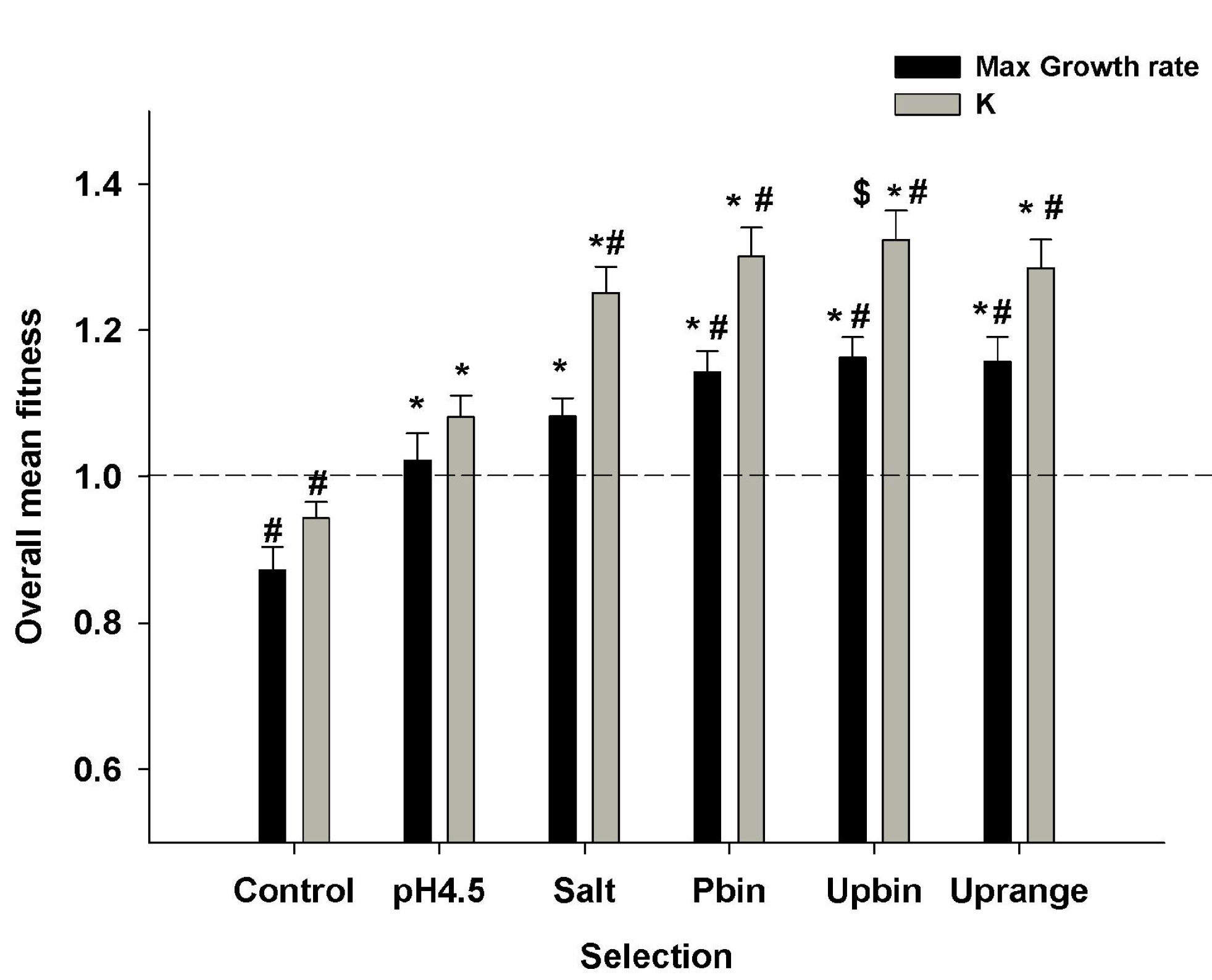
Overall mean (±SE) for fitness. Average fitness for all the selection regimes across three assay environments, namely salt, acid and NB, estimated as maximum growth rate (Black bars) and *K* (Grey bars). Dotted line denotes the fitness of the ancestor. * denotes *p*<0.05 for Tukey HSD when compared with populations selected in pH 4.5 while $ denotes p < 0.05 for Tukey HSD when compared with populations selected in salt.

Since three of the five selection regimes faced multiple environments, it was important to study the variation in fitness across these environments. In order to investigate this, for each selection regime, we calculated the coefficient of variation of fitness for both maximum growth rate and *K* across the 2, *F_5, 114_ assay environments*. All the selected populations had significantly lower CV than the control populations selected in NB for both measures of fitness (Fig 2, *F_5, 114_*= 31.19, *p* < 0.001 for maximum growth rate, *F_5, 114_* = 76.95, *p* < 0.001 for *K*, Table S4 and S5). Previous studies have shown that increased overall mean fitness, after selection in fluctuating environments, is typically accompanied by decreased variation for fitness (reviewed in [25]). Our results were consistent with this observation.

Interestingly, populations selected in constant pH 4.5 environment showed significantly lower *K* (Fig 1) compared to the other four selection regimes (i.e. salt, Pbin, Upbin and Uprange). Similarly, the maximum growth rate of the pH 4.5 selected populations was significantly lower than the three fluctuating regimes (i.e. Pbin, Upbin and Uprange) but not salt and control. Moreover, populations selected in pH4.5 had significantly higher CV than the other four selection regimes (i.e. salt, Pbin, Upbin and Uprange) for both maximum growth rate and *K* (Fig 2). Together, these results suggested that although selection under constant and fluctuating environments resulted in adaptation to all environments containing salt and acid (i.e. everything except Control in Fig 1), the extents of adaptation observed in salt and acid environments were different. Significant statistical interaction between selection and assay environments (*F_10, 342_* = 6.95, *p* < 0.001 for maximum growth rate and *F_10, 342_* = 25.38, *p* < 0.001 for *K*), in conjunction with the results of the post-hoc comparisons (Table S5), showed that the extent of adaptation was greater for all those cases where salt was part of the selection environment (i.e. everything except Control and pH4.5 in Fig 1).

**Fig 2.**
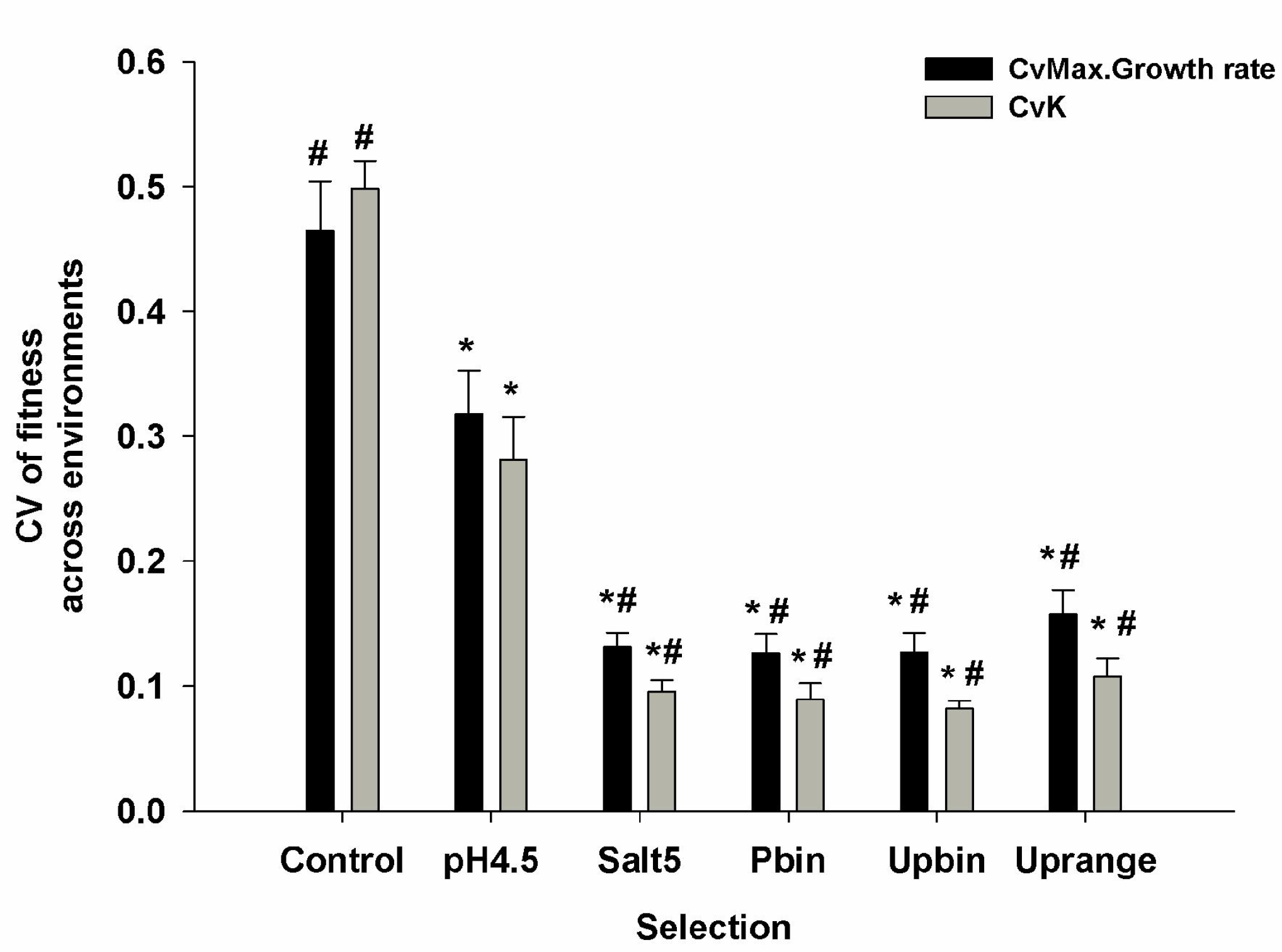
Coefficient of variation (±SE) in fitness. Coefficient of variance across three assay environments, namely salt, acid and NB, was computed for all the selection regimes for maximum growth rate (Black bars) and *K* (Grey bars). * denotes *p* < 0.05 for Tukey HSD when compared to control populations while, # denotes *p* < 0.05 for Tukey HSD when compared with populations selected in pH 4.5.

To confirm this, we performed individual ANOVAs for each of the assay environments. When assayed in salt, for both measures of fitness, selection regime had a significant effect (*F_5, 114_* = 9.19, *p* < 0.001 for maximum growth rate and *F_5, 114_* = 25.2, *p* < 0.001 for *K*). Post-hoc tests (Tukey’s HSD) revealed that in salt, the mean fitnesses were comparable for Pbin, Upbin, Uprange and salt selected populations. While populations selected in acid alone, had significantly lower mean fitness than the rest (Fig 3, Table S7). All the five selected populations showed higher fitness than the control populations (Table S7). When assayed at pH 4.5, the mean fitness did not differ significantly across the selection regimes (*F_5, 114_* = 2.24, *p* = 0.18 for maximum growth rate and *F_5, 114_* = 1.6, p = 0.27 for *K*). When assayed in NB, pH 4.5 selected populations showed significantly lower maximum growth rate than all the other selection regimes and control populations (*F_5, 114_* = 15.88, *p* < 0.001, Table S7, Fig 3) while *K* did not differ significantly across selection regimes (*F_5, 114_* = 2.09, *p* = 0.13). These results confirmed that the differences in the overall mean fitness were primarily driven by the fitnesses of the selected populations under the salt assay environment.

**Fig 3.**
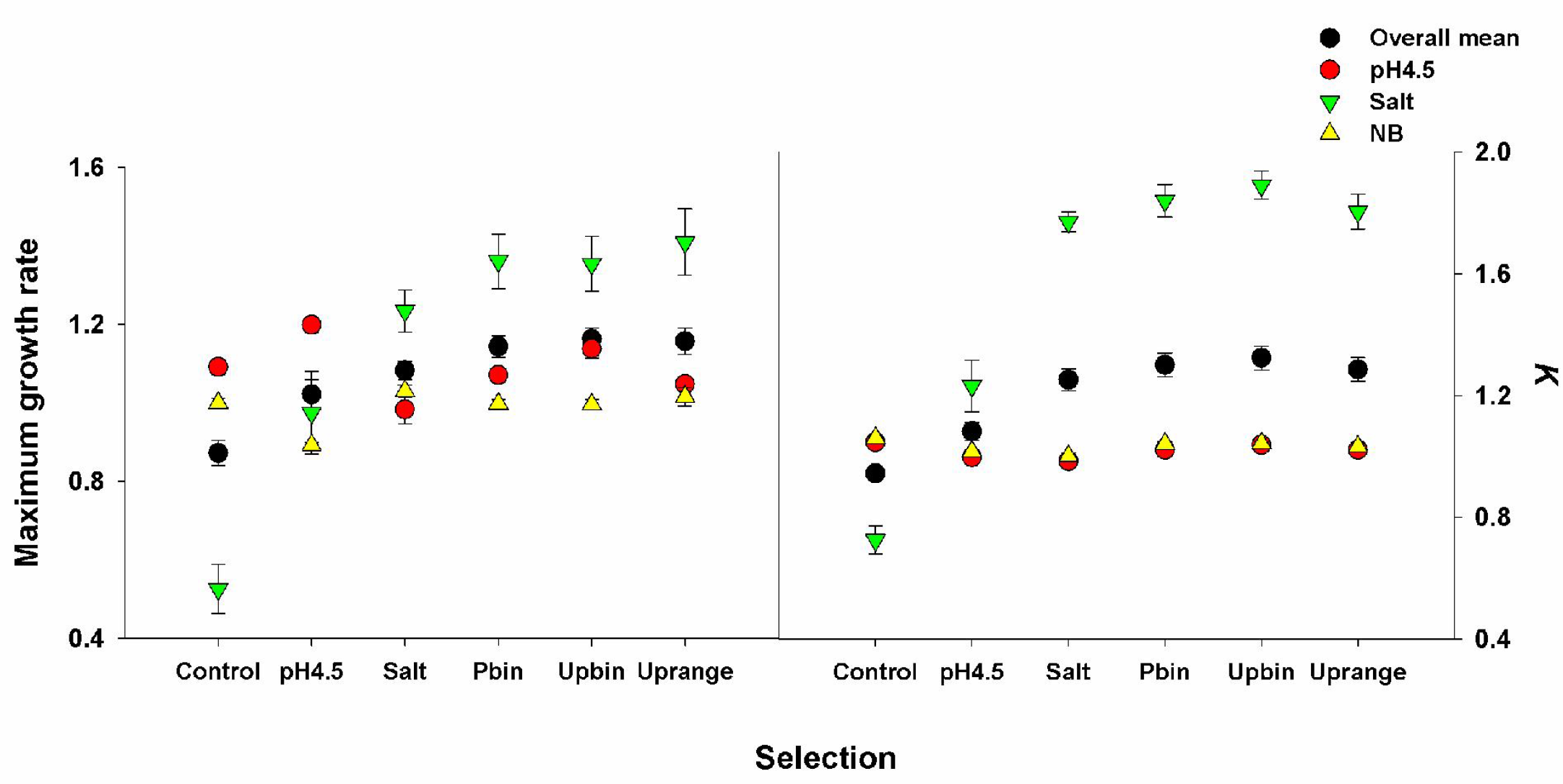
Mean (±SE) fitness. Average fitness for all the selection regimes for (A) Acid (B) Salt (C) NB assay environments, estimated as maximum growth rate (First panel) and *K* (Second panel). Overall mean fitness values, from Fig 1, have been plotted again for easy comparison.

Finally, we analyzed the three fluctuating treatments separately to uncover any subtle differences that might have been lost while analyzing the whole data together. Consistent with the previous analysis, we did not find any difference across the three fluctuating selection regimes, for mean fitness (*F_2, 171_* = 0.094, *p* = 0.91 for maximum growth rate and *F_2, 171_* = 0.44, *p* = 0.68 for *K*, Fig S1). This observation was surprising given that previous studies have shown that extent of adaptation can be different across predictable and unpredictable fluctuations [3, 16] or even across different kinds of unpredictable fluctuations [3].

## Discussion

Results of previous experimental evolution studies have led to important insights about the fitness outcomes of selection under fluctuating environments [25, 26]. For instance, when subjected to predictable fluctuations, organisms improve fitness over the entire range of environments experienced [17, 18, 27]. On the other hand, the outcomes of selection under unpredictable fluctuations are equivocal as the fitness can increase [18, 19], show no change [3] or increase in some selection environments but not in others [16]. While these studies have focused on the effects of a single stress at a time, other investigations have looked at the outcomes of selection in the presence of multiple stresses [12, 13]. When a population experiences multiple unpredictably fluctuating stresses simultaneously, then fitness can be improved over the long term (~ 900 generations) [13] but not over the short term (~170 generations) [12]. This leads one to question the role of unpredictability in this context. In other words, is it the unpredictability of the fluctuations or the fact that multiple stresses are acting simultaneously, or an interaction of the two, that prevents the adaptation in short run? Here we show that the extent of increase in overall mean fitness is comparable for predictable and unpredictable fluctuations over short run when complexity is absent (Fig 1). This suggests that the complexity of the environment, i.e. simultaneous presence of multiple environmental factors, may pose a bigger constraint on extent of adaptation than unpredictability of fluctuations, at least over short time scales and for the chosen environmental parameters. Our results are also in contrast with previous studies comparing the fitness effects of predictable and unpredictable fluctuations on multiple values of a single parameter like temperature [3, 27] or pH [16], which have reported reduced adaptation for unpredictable fluctuations. We observe that when fluctuations occur across qualitatively different environments (like salt and acid), extent of adaptation is comparable across predictable and unpredictable selection regimes (Fig 1).

Our results also uncover another major factor governing the extent of adaptation, which is the choice of the selection environment, or more specifically, the extent of ancestral adaptation to it. The fitness improvement when assayed in salt environment explains most of the increase observed in overall mean fitness (Fig 3). Not surprisingly, populations selected in salt alone, show overall mean fitness comparable to populations selected in fluctuating regimes (Fig1). Concurrently, populations selected in pH 4.5 do not fare better than the populations selected in NB (Fig1). This result is not surprising given that *E. coli* is known to be well adapted to acidic environments (reviewed in [28]). These observations together suggest that when there are fluctuations across multiple environmental parameters, extent of ancestral adaptation to individual environments will play a key role in determining the magnitude of adaptation. When the ancestor is poorly adapted to some of the components of the selection environments (here salt), adaptation will mainly be driven by this single factor.

Comparison across studies that deal with constant selection environments allows us to determine the role played by the choice of environment in the context of the model system used. Experiments involving fluctuating environments, on the other hand, pose a major logistic hurdle in terms of the relevant controls involved in this context. It is nearly impossible to study all three axes of fluctuations, namely predictability, complexity and grain size, together along with the controls for every selection environment chosen. Use of different model systems and environmental parameters makes it even more difficult to compare outcomes across studies. As a result, experimental investigations involving same model system and environmental parameters can provide important insights Results of this study, together with the previous work done with the suggests that the extent of adaptation of ancestor will strongly affect the fitness outcomes [12, 13]. The nature of fluctuations themselves, whether predictable or unpredictable, might be of secondary importance when ancestor is poorly adapted to any of the selection environments.

All else being equal, this is a reason for cautious optimism for anyone who is worried about the effects of increasing climatic variability [1, 2] on the evolution of microbes [12]. These results suggest that at least the unpredictability component of environmental fluctuations might be relatively inconsequential in determining microbial evolution in the short term. However, all else is seldom equal in biology and therefore, due caution must be exercised in extrapolating this result to other bacteria or other kinds of environmental conditions. But one can definitely state that the intricacies of the relationships between various components of environmental fluctuations in shaping the evolutionary dynamics of organisms will be a major challenge for evolutionary biologists for at least some time in the future.

## Acknowledgements

SK was supported by a Senior Research Fellowship from Council of Scientific and Industrial Research, Govt. of India. This project was supported by grant# BT/PR5655/BRB/10/1088/2012 from Department of Biotechnology, Government of India and internal funding from Indian Institute of Science Education and Research, Pune. The authors declare no conflict of interests.

---

**S1. Details of the ancestral *Escherichia coli* population used for the selection**

We used an *Escherichia coli* K12 MG1655 strain in which the lacY gene had been replaced with a Kanamycin resistance gene. Colonies of this bacterium are white coloured on MacConkey’s agar as opposed to the red coloured colonies produced by other *Escherichia coli*.

**S2. Composition of Nutrient broth -**

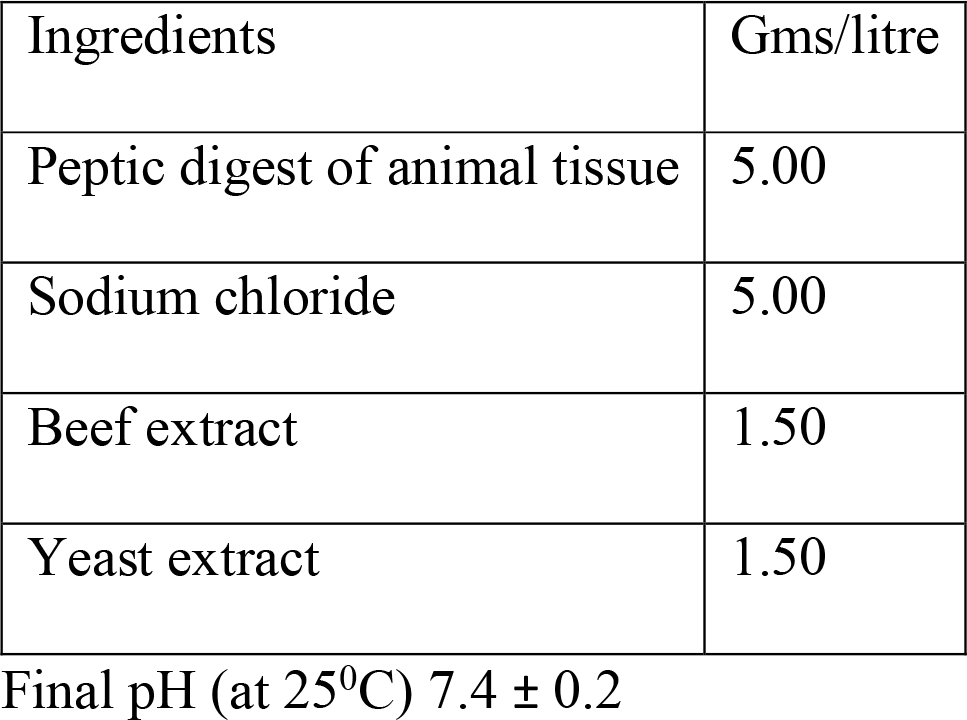

For making NB^Kan^, 0.05 mg/ml of Kanamycin was added to the above mixture after autoclaving and cooling.

We added 2g/100ml of agar to the above mixture before autoclaving, to make Nutrient Agar^Kan^.

**S3.** Composition of three fluctuating selection regimes Pbin (predictable binary), Upbin (unpredictable binary) and Uprange (unpredictable range)

| **Day** | **Pbin** | **Upbin** | **Uprange** |
| --- | --- | --- | --- |
| 1 | pH 4.5 | pH 4.5 | pH 4.5 |
| 2 | Salt 5 g% | pH 4.5 | Salt 5 g% |
| 3 | pH 4.5 | Salt 5 g% | Salt 3.5 g% |
| 4 | Salt 5 g% | pH 4.5 | pH 4.5 |
| 5 | pH 4.5 | Salt 5 g% | pH 4.5 |
| 6 | Salt 5 g% | Salt 5 g% | pH 4.5 |
| 7 | pH 4.5 | pH 4.5 | pH 4.5 |
| 8 | Salt 5 g% | pH 4.5 | pH 4.5 |
| 9 | pH 4.5 | pH 4.5 | Salt 5 g% |
| 10 | Salt 5 g% | pH 4.5 | pH 4.5 |
| 11 | pH 4.5 | Salt 5 g% | Salt 4 g% |
| 12 | Salt 5 g% | pH 4.5 | Salt 4 g% |
| 13 | pH 4.5 | Salt 5 g% | Salt 3.5 g% |
| 14 | Salt 5 g% | pH 4.5 | Salt 5 g% |
| 15 | pH 4.5 | pH 4.5 | pH 4.5 |
| 16 | Salt 5 g% | pH 4.5 | Salt 4.5 g% |
| 17 | pH 4.5 | pH 4.5 | Salt 4 g% |
| 18 | Salt 5 g% | pH 4.5 | Salt 3.5 g% |
| 19 | pH 4.5 | Salt 5 g% | Salt 4 g% |
| 20 | Salt 5 g% | pH 4.5 | pH 4.5 |
| 21 | pH 4.5 | pH 4.5 | Salt 3 g% |
| 22 | Salt 5 g% | pH 4.5 | Salt 3 g% |
| 23 | pH 4.5 | pH 4.5 | Salt 4 g% |
| 24 | Salt 5 g% | Salt 5 g% | Salt 2 g% |
| 25 | pH 4.5 | Salt 5 g% | Salt 3.5 g% |
| 26 | Salt 5 g% | pH 4.5 | Salt 2 g% |
| 27 | pH 4.5 | Salt 5 g% | Salt 4 g% |
| 28 | Salt 5 g% | pH 4.5 | Salt 2 g% |
| 29 | pH 4.5 | Salt 5 g% | Salt 3.5 g% |
| 30 | Salt 5 g% | pH 4.5 | Salt 4.5 g% |

**Table S4.** Overall mean fitness, CV for overall mean fitness and mean fitness in individual selection environments, where fitness is measured as maximum growth rate and K, for all the selection regimes.

| Assay | Selection regime | Mean fitness (maximum growth rate) ±SEM | Mean fitness (*K*) ±SEM |
| --- | --- | --- | --- |
| **Overall mean fitness** | Control | 0.87±0.03 | 0.94±0.02 |
| | Pbin | 1.14±0.02 | 1.30±0.03 |
| | Upbin | 1.16±0.02 | 1.32±0.04 |
| | Uprange | 1.15±0.03 | 1.28±0.03 |
| | pH4.5 | 1.02±0.03 | 1.08±0.02 |
| | Salt | 1.08±0.02 | 1.25±0.03 |
| **CV for overall mean fitness** | Control | 0.46±0.03 | 0.49±0.02 |
| | Pbin | 0.12±0.01 | 0.08±0.01 |
| | Upbin | 0.12±0.01 | 0.08±0.006 |
| | Uprange | 0.15±0.01 | 0.10±0.01 |
| | pH4.5 | 0.31±0.03 | 0.28±0.03 |
| | Salt | 0.13±0.01 | 0.09±0.009 |
| **Mean fitness in pH4.5** | Control | 1.09±0.01 | 1.04±0.008 |
| | Pbin | 1.07±0.01 | 1.02±0.009 |
| | Upbin | 1.13±0.02 | 1.03±0.01 |
| | Uprange | 1.04±0.01 | 1.02±0.007 |
| | pH4.5 | 1.19±0.01 | 0.99±0.009 |
| | Salt | 0.98±0.03 | 0.98±0.01 |
| **Mean fitness in Salt** | Control | 0.52±0.06 | 0.72±0.04 |
| | Pbin | 1.36±0.06 | 1.84±0.05 |
| | Upbin | 1.35±0.06 | 1.89±0.04 |
| | Uprange | 1.40±0.08 | 1.80±0.05 |
| | pH4.5 | 0.97±0.10 | 1.23±0.08 |
| | Salt | 1.23±0.05 | 1.77±0.03 |
| **Mean fitness in NB** | Control | 0.99±0.01 | 1.06±0.007 |
| | Pbin | 0.99±0.009 | 1.04±0.006 |
| | Upbin | 0.99±0.01 | 1.04±0.006 |
| | Uprange | 1.01±0.02 | 1.03±0.01 |
| | pH4.5 | 0.89±0.006 | 1.01±0.006 |
| | Salt | 1.03±0.01 | 1.00±0.01 |

**Table S5.** Tukey for overall mean and coefficient of variance for fitness, measured as maximum growth rate and *K*. *p* < 0.05 denotes significant difference. Significant values have been italicized for convenience.

| **Overall mean maximum growth rate** |  |  |  |  |  |  |
| --- | --- | --- | --- | --- | --- | --- |
|  | Control | Pbin | Upbin | Uprange | pH4.5 | Salt |
| Control |  | *0.000020* | *0.000020* | *0.000020* | *0.000061* | *0.000020* |
| Pbin | *0.000020* |  | 0.990786 | 0.997929 | *0.001984* | 0.400680 |
| Upbin | *0.000020* | 0.990786 |  | 0.999984 | *0.000167* | 0.122168 |
| Uprange | *0.000020* | 0.997929 | 0.999984 |  | *0.000330* | 0.177301 |
| pH4.5 | *0.000061* | *0.001984* | *0.000167* | *0.000330* |  | 0.401039 |
| Salt | *0.000020* | 0.400680 | 0.122168 | 0.177301 | 0.401039 |  |
| **Overall mean $K$** |  |  |  |  |  |  |
|  | Control | Pbin | Upbin | Uprange | pH4.5 | Salt |
| Control |  | *0.000020* | *0.000020* | *0.000020* | *0.000020* | *0.000020* |
| Pbin | *0.000020* |  | 0.923008 | 0.986137 | *0.000020* | 0.267850 |
| Upbin | *0.000020* | 0.923008 |  | 0.568433 | *0.000020* | *0.022164* |
| Uprange | *0.000020* | 0.986137 | 0.568433 |  | *0.000020* | 0.679018 |
| pH4.5 | *0.000020* | *0.000020* | *0.000020* | *0.000020* |  | *0.000020* |
| Salt | *0.000020* | 0.267850 | *0.022164* | 0.679018 | *0.000020* |  |
| **CV for Maximum growth rate** |  |  |  |  |  |  |
|  | Control | Pbin | pH4.5 | Salt | Upbin | Uprange |
| Control |  | *0.000119* | *0.001050* | *0.000119* | *0.000119* | *0.000119* |
| Pbin | *0.000119* |  | *0.000124* | 0.999992 | 1.000000 | 0.952372 |
| pH4.5 | *0.001050* | *0.000124* |  | *0.000128* | *0.000124* | *0.000344* |
| Salt | *0.000119* | 0.999992 | *0.000128* |  | 0.999996 | 0.977968 |
| Upbin | *0.000119* | 1.000000 | *0.000124* | 0.999996 |  | 0.955966 |
| Uprange | *0.000119* | 0.952372 | *0.000344* | 0.977968 | 0.955966 |  |
| **CV for $K$** |  |  |  |  |  |  |
|  | Control | Pbin | pH4.5 | Salt | Upbin | Uprange |
| Control |  | *0.000119* | *0.000119* | *0.000119* | *0.000119* | *0.000119* |
| Pbin | *0.000119* |  | *0.000119* | 0.999892 | 0.999827 | 0.983811 |
| pH4.5 | *0.000119* | *0.000119* |  | *0.000119* | *0.000119* | *0.000119* |
| Salt | *0.000119* | 0.999892 | *0.000119* |  | 0.995645 | 0.997995 |
| Upbin | *0.000119* | 0.999827 | *0.000119* | 0.995645 |  | 0.932573 |
| Uprange | *0.000119* | 0.983811 | *0.000119* | 0.997995 | 0.932573 |  |

**Table S6.**
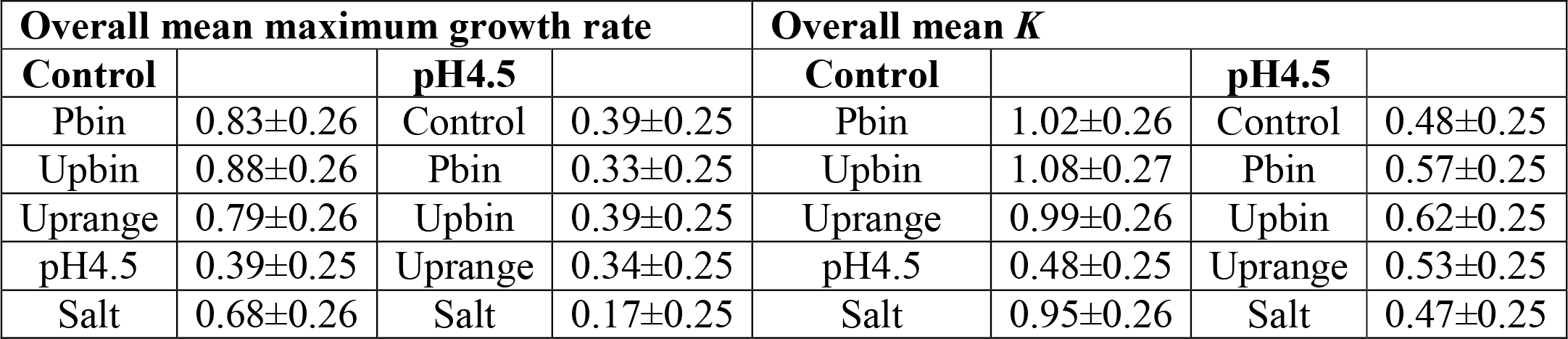
Effect size comparisons of control populations and populations selected in pH 4.5 with all the other selection treatments for both measures of fitness, overall mean maximum growth rate and *K*. Effect size is measured as Cohen’s d (± 95%CI) (REF) and interpreted as small, medium and large for 0.2 < d < 0.5, 0.5 < d < 0.8 and d > 0.8, respectively.

**Table S7.** Tukey for mean fitness for the cases where ANOVA showed significant effect of selection. These include Tukey values for fitness, measured as maximum growth rate and *K*, when assayed in salt. Tukey for mean fitness, measured as maximum growth rate in NB. *p* < 0.05 denotes significant difference. Significant values have been italicized for convenience.

| **Maximum growth rate in Salt** |  |  |  |  |  |  |
| --- | --- | --- | --- | --- | --- | --- |
|  | Control | Pbin | Upbin | Uprange | pH4.5 | Salt |
| Control |  | *0.000119* | *0.000119* | *0.000119* | *0.000144* | *0.000119* |
| Pbin | *0.000119* |  | 1.000000 | 0.994524 | *0.000600* | 0.721176 |
| Upbin | *0.000119* | 1.000000 |  | 0.989832 | *0.000761* | 0.766058 |
| Uprange | *0.000119* | 0.994524 | 0.989832 |  | *0.000176* | 0.378949 |
| pH4.5 | *0.000144* | *0.000600* | *0.000761* | *0.000176* |  | *0.049994* |
| Salt5 | *0.000119* | 0.721176 | 0.766058 | 0.378949 | *0.049994* |  |
| **K in Salt** |  |  |  |  |  |  |
|  | Control | Pbin | Upbin | Uprange | pH4.5 | Salt |
| Control |  | *0.000119* | *0.000119* | *0.000119* | *0.000144* | *0.000119* |
| Pbin | *0.000119* |  | 1.000000 | 0.994524 | *0.000600* | 0.721176 |
| Upbin | *0.000119* | 1.000000 |  | 0.989832 | *0.000761* | 0.766058 |
| Uprange | *0.000119* | 0.994524 | 0.989832 |  | *0.000176* | 0.378949 |
| pH4.5 | *0.000144* | *0.000600* | *0.000761* | *0.000176* |  | *0.049994* |
| Salt5 | *0.000119* | 0.721176 | 0.766058 | 0.378949 | *0.049994* |  |
| **Maximum growth rate in NB** |  |  |  |  |  |  |
|  | Control | Pbin | Upbin | Uprange | pH4.5 | Salt |
| Control |  | 1.000000 | 0.999949 | 0.958238 | *0.000120* | 0.578978 |
| Pbin | 1.000000 |  | 0.999996 | 0.938905 | *0.000120* | 0.525733 |
| Upbin | 0.999949 | 0.999996 |  | 0.897844 | *0.000121* | 0.443231 |
| Uprange | 0.958238 | 0.938905 | 0.897844 |  | *0.000119* | 0.969284 |
| pH4.5 | *0.000120* | *0.000120* | *0.000121* | *0.000119* |  | *0.000119* |
| Salt5 | 0.578978 | 0.525733 | 0.443231 | 0.969284 | *0.000119* |  |

**Fig S1.**
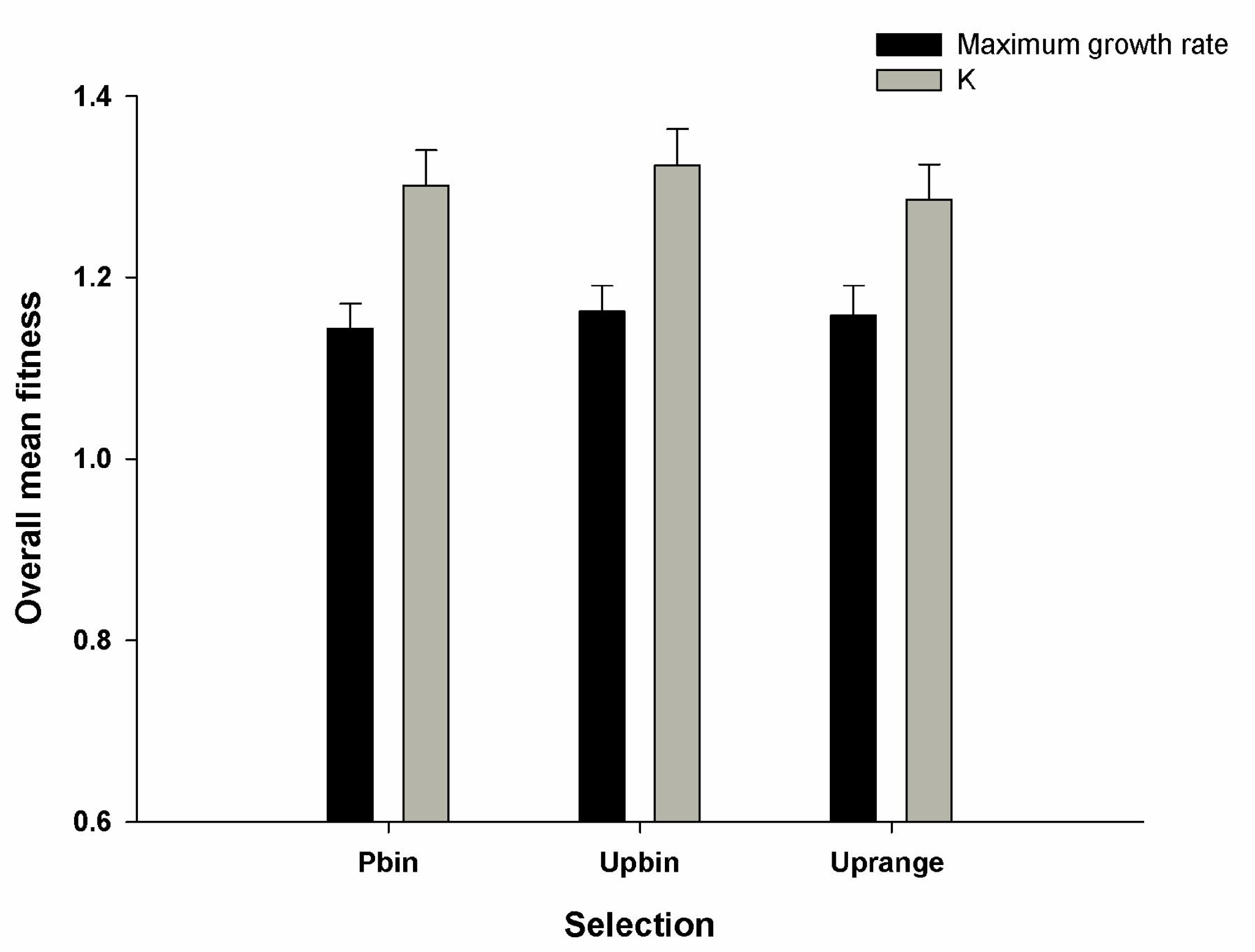
Overall mean (±SE) for fitness. Average fitness for three fluctuating selection regimes across three assay environments, namely salt, acid and NB, estimated as maximum growth rate (Black bars) and *K* (Grey bars).

